# Direct Small Molecule Activation of Mitofusins

**DOI:** 10.1101/301713

**Authors:** Emmanouil Zacharioudakis, Nikolaos Biris, Thomas P. Garner, Yun Chen, Ryan Pekson, Rimpy Dhingra, Gaetano Santulli, Lorrie A. Kirshenbaum, Richard N. Kitsis, Evripidis Gavathiotis

## Abstract

Mitochondrial fusion is a physiological process that is regulated by mitofusins on the outer mitochondrial membrane. Conformational plasticity between anti- and pro-tethering conformations of mitofusins permits mitochondrial tethering and subsequent fusion. Here we developed a pharmacophore-based model to rationally manipulate the conformational plasticity of mitofusin 2 and perfomed an *in silico* small-molecule screen. This enabled the discovery of a direct activator of mitofusins, MASM7, capable of potently promoting mitochondrial fusion. The specificity of the MASM7-mitofusin 2 interaction is highlighted by structure-activity relationships of MASM7 analogues, FRET, NMR and mitochondrial fusion studies using mitofusin mutants. Our study identified the first-in-class direct activator of mitofusins, demonstrating a new paradigm for chemical modulation of mitochondrial fusion and downstream processes.

## Introduction

Mitochondria are highly dynamic organelles that fuse and divide constantly. The physiological processes that regulate this mitochondrial dynamism are fusion and fission, respectively (Detmer *et al.*, 2007). A large variability in mitochondrial morphology is observed among different cell types and within the same cell, as mitochondria can morph into small spheres, short rods or long tubules. This dynamism allows mitochondria to exchange contents (e.g. lipid membranes, proteins), promote repair, and maintain mitochondrial quality control (Shirihai *et al.*, 2015).

The fusion machinery includes mitofusin (Mfn) 1 and 2 on the outer mitochondrial membrane (OMM) and optic atrophy (Opa) 1 on the inner membrane (Shirihai *et al.*, 2015; Schrepfer *et al.*, 2016). Mfn1 and Mfn2 share high sequence homology (Schrepfer *et al*., 2016). Both homologs possess an N-terminal GTPase domain, a coiled-coiled heptad repeat (HR1) domain, a short transmembrane domain responsible for anchoring Mfns on the OMM, and a second coiled-coiled heptad repeat (HR2) domain located in the C-terminus (Figure 1A) (Shirihai *et al.*, 2015; Schrepfer *et al.*, 2016; Koshiba *et al.*, 2004). The initial step in fusion is tethering of the OMMs of adjacent mitochondria. HR2 domains of Mfns from two adjacent mitochondria interact in an anti-parallel trans manner to form homotypic (Mfn1-Mfn1 or Mfn2-Mfn2) or heterotypic (Mfn1-Mfn2) complexes, subsequently mediating mitochondrial tethering (Koshiba *et al.*, 2004). We recently proposed a model in which Mfn2 can undergo conformational activation to promote mitochondrial tethering (Franco *et al.*, 2016). Based on this model, Mfn2 can adopt anti- or pro-tethering conformations (Figure 1A) (Franco *et al.*, 2016). In the anti-tethering conformation HR2 interacts intra-molecularly with HR1 in an anti-parallel manner (Figure 1A). On the other hand, in the pro-tethering conformation HR1-HR2 interactions are disrupted, and the HR2 domain extends into the cytosol to mediate mitochondrial tethering through anti-parallel HR2-HR2 interactions as proposed by Koshiba *et al.* (2004) (Figure 1A). Recently, Cao *et al.* (2017) demonstrated that dimerization of GTPase domains of mitofusins may also control mitochondrial tethering. In contrast, Mattie *et al.* (2018) proposed an alternative model for mitochondrial tethering, in which Mfns are activated by HR2 domain dimerization in the intermembrane mitochondrial space. Further work will be needed to resolve the differences in these models.

**Figure 1.**
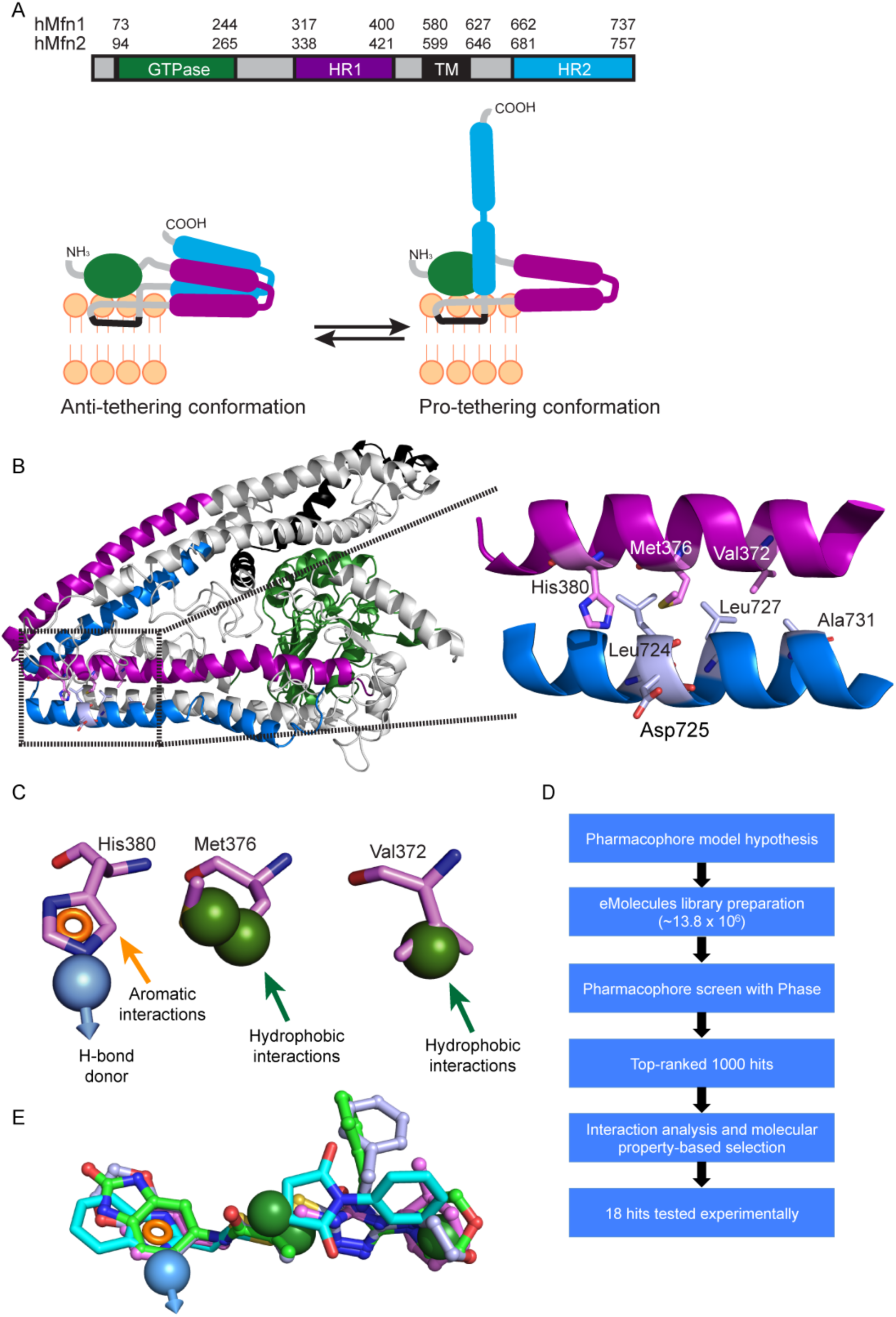
Pharmacophore model-based strategy for *in silico* discovery of MASMs. (A) Mfn2 undergoes conformational changes to promote mitochondrial tethering. Anti- and pro-tethering conformations of Mfn2. (B) Ribbon representation of the human Mfn2 structural model and HR1-HR2 interactions. (C) Pharmacophore hypothesis based on the interactions of the side chains of the HR1-amino acids: Val372, Met376, His380. (D) Schematic representation of a *in silico*-based screening strategy for the discovery of putative MASMs. (E) A set of small molecules shown in different color demonstrates how selected molecules fulfill the criteria of the pharmacophore hypothesis.

Irrespective of the precise tethering model, however, we and others have shown disruption of the HR1-HR2 interaction is critical for mitochondrial tethering and fusion to take place (Franco *et al.* 2016, Huang *et al.*, 2011). Therefore, we sought to identify mitofusin activating small molecules (MASMs) by targeting the disruption of the HR1-HR2 interaction. To accomplish this, we used the HR1-HR2 interaction that is supported by the structural model of full-length Mfn2 (Franco *et al.* 2016) and the crystal structures of Mfn1 (Cao *et al.*, 2017).

Derangements/imbalances in mitochondrial dynamics can lead to neurological, metabolic, cardiovascular disorders, as well as cancer (Detmer *et al.*, 2007; Shirihai *et al.*, 2015; Schrepfer *et al.*, 2016). For example, impaired mitochondrial fusion in the rare neurological disease Charcot Marie Tooth disease type 2A (CMT2A), can result from a plethora of loss of function mutations in Mfn2 (Bombelli *et al.*, 2014). Indeed, we have shown that the HR1-based peptide, 367-384 Gly and the small molecule described herein can correct the CMT2A Mfn2 mutant phenotype (Franco *et al.*, 2016; Rocha *et al.*, 2018). This paper describes the discovery of this small molecule, which may provide a drug prototype for CMT2A as well as for more common acquired disorders mediated by defects in mitochondrial fusion.

## RESULTS AND DISCUSSION

To identify small molecules that directly activate Mfn2 and promote mitochondrial fusion, we subjected Mfn2 to a structure-guided pharmacophore-based approach combined with *in silico* screening with the goal of directly promoting the pro-tethering and/or inhibiting the formation of anti-tethering conformation of Mfn2. Our previous structural model of full-length Mfn2 in the anti-tethering conformation (Franco *et al.*, 2016) and the recent crystal structure of Mfn1 (Cao, *et al.*, 2017) informed us about the intra-molecular HR1-HR2 interactions,. Visual inspection of the intra-molecular HR1-HR2 interactions in the human Mfn2 structural model and particularly in the region that involves HR1-residues: 367-384, which our previous work using a 367-384 Gly peptide suggested to regulate, provided structural insights for small molecule mimicry (Franco *et al.*, 2016). Specifically, several hydrophobic interactions were observed between the following HR1-amino acids: Val372, Met376 and HR2-amino acids: Leu724, Leu727 and Ala731 and a possible hydrogen bond between the HR1-amino acid: His380 and HR2-amino acid: Asp725 (Figure 1B). All of these residues are highly conserved in Mfn2 and Mfn1 of different species (Figure S1A). We envisioned that a small molecule capable of recapitulating the interactions of the aforementioned HR1-amino acids would compete with intra-molecular HR1-HR2 interactions and promote Mfn2 conformational activation. Accordingly, to identify such a small molecule, we generated a pharmacophore hypothesis that comprises the following features: three hydrophobic interactions, an aromatic interaction and a hydrogen bond donor based on the interactions of the HR1 residues: Val372, Met376 and His380 (Figure 1C). An *in silico* library of 13.8 × 10^6^ commercially available small molecules was screened to fit the pharmacophore hypothesis using PHASE (Figure 1D) (Dixon *et al.*, 2006). The top 1000 hits of the *in silico* screen were clustered for diversity and analyzed based on the fit to residues of the HR1 domain, their interactions with residues of the HR2 domain, and their molecular properties. Moreover, a number of filters (e.g. elimination of hits with poor ADMET properties using Qikprop) were applied to provide hits with drug-like properties (Duffy *et al.*, 2000). Finally, a subset of 18 compounds was selected for experimental validation based on their fit to the pharmacophore hypothesis and molecular diversity of their scaffolds (Figures 1D, 1E).

To evaluate the capacity of the selected hits to promote a conformational activation of Mfn2, a cell-based screening assay was developed based on a FRET biosensor of Mfn2, a molecular tool that we had previously used to monitor the conformational changes of Mfn2 in cells (Franco *et al.*, 2016). Based on the structural model of human Mfn2 in the anti-tethering conformation and in agreement with the crystal structure of Mfn1, the N-terminus is in approximate distance with the C-terminus in the HR2 domain. Therefore the N-terminus was fused with mCerulean and the C-terminus in the HR2 domain was fused with mVenus (Figure 2A). This FRET-based Mfn2 biosensor was transiently transfected in HEK 293T cells and co-localized with ATP synthase subunit beta (ATP5B), consistent with mitochondrial localization (Figure S1B). Loss of intra-molecular HR1-HR2 interactions leads to a conformational change and promotes the pro-tethering conformation of Mfn2. The increased distance between the N-terminus and the C-terminus, occurring in the pro-tethering conformation of Mfn2, is expected to lead to a decrease in the resonance energy that is transferred (Figure 2A). Indeed, we previously showed that the HR1-derived peptide: 367-384 Gly that promotes the pro-tethering conformation of Mfn2 was able to decrease FRET, whereas the HR1-derived peptide: 398-418 Gly that promotes the anti-tethering conformation of Mfn2 was able to increase FRET (Franco *et al.*, 2016). Therefore, we used the HR1-derived peptide: 367-384 Gly as a positive control in our assay. Next, we examined the ability of the 18 putative MASMs to promote the pro-tethering conformation of Mfn2, using a decrease in FRET as a readout (Figure 2B, Table S1). Strikingly, MASM7 was the only small molecule in our screen that significantly decreased FRET at 10 μM, indicative of promoting the pro-tethering conformation of Mfn2 (Figure 2B). MASM7 was resynthesized to document its purity by ^1^H and ^13^C NMR and reconfirm its activity (Figures 2C and S2A).

**Figure 2.**
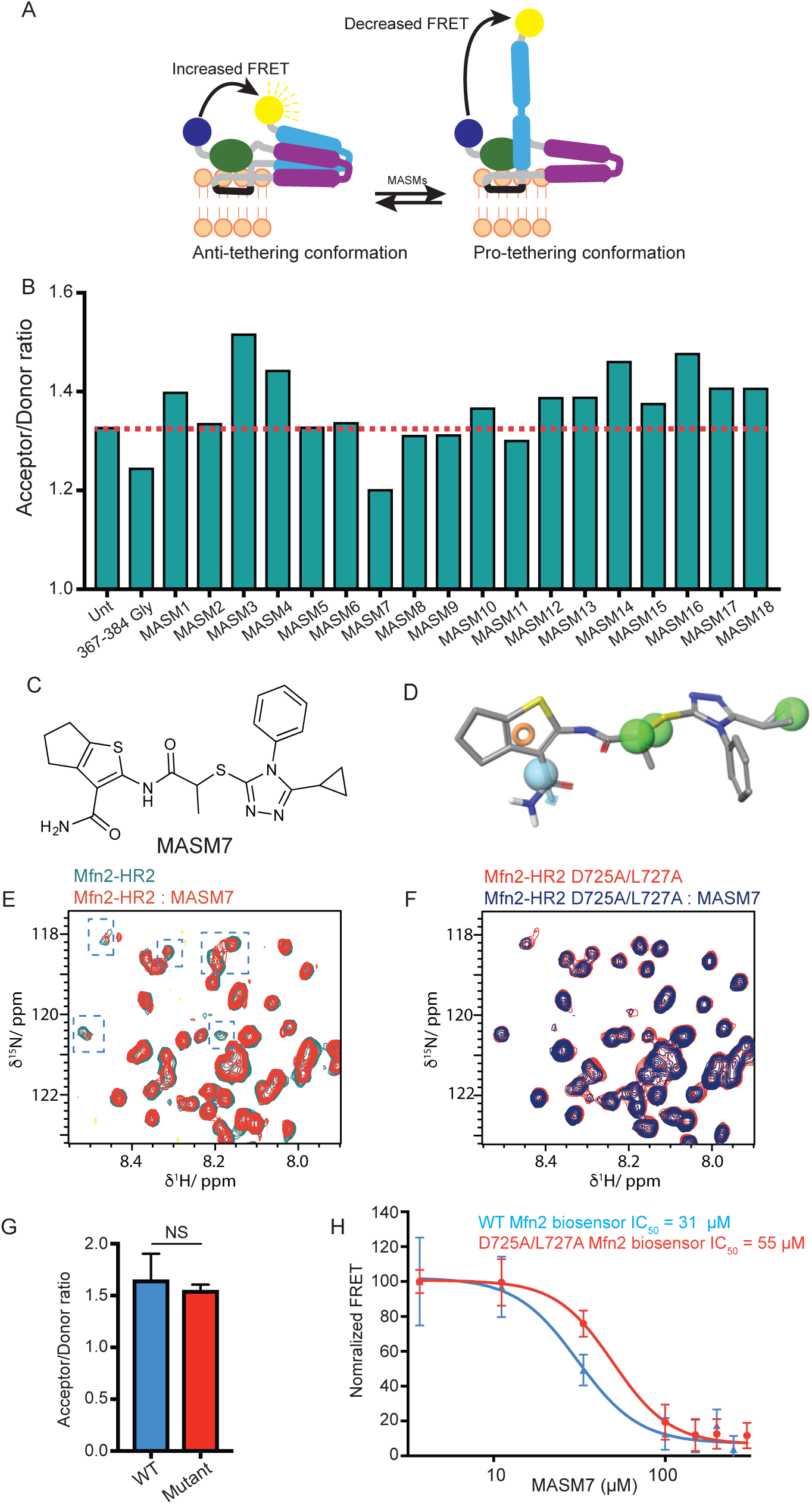
MASM7 promotes the pro-tethering conformation of Mfn2 through binding to the HR2 domain. (A) Schematic representation of FRET-based Mfn2 biosensor conformational changes. Cyan sphere represents mCerulean and yellow sphere represent mVenus. (B) Screening of putative MASMs using FRET-based Mfn2 biosensor in HEK 293T cells. Cells were treated with MASMs or 367-384 Gly (10 μM, 2 h). We used 367-384 Gly peptide as a positive control (Franco *et al.*, 2016). (C) Structure of MASM7. (D) MASM7 satisfies the pharmacophore hypothesis features as described in our methodology. Green spheres represent hydrophobic interactions of the cyclopropane and methyl groups. Cyan sphere represents hydrogen bond interaction of the amide group and the orange ring represents aromatic interaction of the thiophene group. (E) Zoomed region of overlaid ^1^H-^15^N HSQC spectra of Mfn2-HR2 (50 μM) with (red cross-peaks) and without (green cross-peaks) MASM7 (200 μM). Line broadening and chemical shifts upon addition of MASM7 are highlighted with blue boxes. (F) Zoomed region of overlaid ^1^H-^15^N HSQC spectra of Mfn2-HR2 D725A/L727A mutant (50 μM) with (blue crosspeaks) and without (red cross-peaks) MASM7 (200 μM). Effects of MASM7 on broadening and chemical shifts are lost. See full spectra in (Figure S5). (G) Acceptor/Donor ratios of WT and D725A/L727A FRET based Mfn2 biosensors. Error bars represent mean ± SD, *n* = 3. (H) D725A/L727A FRET based Mfn2 biosensor decreased potency of MASM7. Error bars represent mean ± SEM, *n* = 4.

MASM7 possesses functional groups that fulfill the 5 criteria of pharmacophore hypothesis (Figure 2D). Specifically, the substituted thiophene ring mimics the side chain of His380 and the hydrophobic methyl and the cyclopropane groups mimic the side chains of Met376 and Val372, respectively. We evaluated the importance of each functional group using the FRET-based Mfn2 biosensor (Figure S2B). Loss of the thiophene ring diminished significantly the ability of this scaffold to reduce FRET. Moreover, the replacement of the methyl group of MASM7 with hydrogen decreased the potency of the compound. Replacement of the cyclopropane group of MASM7 with bulky substituents (e.g. benzene, methyl-furan) was not tolerated, while the replacement with the furan group was well tolerated. These data suggested that the aforementioned functional groups of MASM7 play a key role in its interactions with HR2 to promote a conformational change on Mfn2.

Previous structural studies reported that the HR2 domain adopts a helical conformation in the anti-tethering and pro-tethering conformations of Mfn2 (Koshiba *et al.*, 2004; Franco *et al.*, 2016; Cao *et al.*, 2017). MASM7 was designed to bind the HR2 domain and may bind to the proposed residues on HR2 domain in either anti- or pro-tethering conformation (Figures 1A and 1B). Therefore, to validate that MASM7 directly interacts with the HR2 domain *in vitro*, we produced for the first time ^15^N-labeled Mfn2-HR2 domain (residues 678-757) and recorded heteronuclear quantum spin correlation (HSQC) spectra with or without MASM7 (Figures 2E and S2C). HSQC NMR spectra of the isolated HR2 domain showed evidence of a folded conformation within HR2. Titration of MASM7 revealed peak broadening corresponding to a number of HR2 residues, demonstrating direct binding to the HR2 domain indicative of a low μM binding affinity (Figures 2E and S2C). On the other hand, titration of MASM7 to an ^15^N-labeled Mfn2-HR2 D725A/L727A, mutations that are predicted to disrupt MASM7 binding (Figure 1B), in fact demonstrated no evidence of MASM7 binding even at 200 μM (Figures 2F and S2D). Thus, these NMR data suggest that MASM7 binds directly to the HR2 domain of Mfn2 and that D725 and L727 residues are critical for this interaction, consistent with our pharmacophore model.

To further validate that MASM7 interacts with the HR2 in cells, we also generated the same double mutant (D725A/L727A) in the FRET-based Mfn2 biosensor. Wild type (WT) and D725A/L727A FRET-based Mfn2 biosensors presented similar acceptor/donor ratios, suggesting that these double point mutations did not alter structural transformation of Mfn2 between pro- and anti-tethering conformations (Figure 2G). Titration of MASM7 reduced FRET in a concentration-dependent manner. Of note, the D725A/L727A Mfn2 biosensor required almost twice the concentration of MASM7 (IC_50_ = 55 μM) for the reduction of FRET compared to the WT biosensor (IC_50_ = 31 μM) (Figure 2H). Therefore, our results further validated the interaction of MASM7 with D725 and L727 on the HR2 domain of Mfn2 and the promotion of pro-tethering conformation by MASM7.

Next, we examined if the pro-tethering activity of MASM7 can also trigger mitochondrial fusion in cells. We examined whether MASM7 could induce mitochondrial fusion in wild type MEFs and MEFs lacking Mfn1, Mfn2, or both, using mitochondrial aspect ratio (mitochondrial length/width, reflecting fusion) as a readout. Notably, MASM7 increased significantly mitochondrial aspect ratio in WT MEFs. The same was observed, albeit to a lesser degree, in MFN1 KO and MFN2 KO MEFs (Figures 3A, B and C). In contrast, MASM7 did not increase mitochondrial aspect ratio in MFN1/MFN2 double knockout (DKO) MEFs. Collectively, these data indicate that MASM7 can induce mitochondrial fusion by directly activating either Mfn2 or Mfn1 and MASM7-induction of mitochondrial fusion is greater when both Mfn2 and Mfn1 are expressed. This result is in agreement with the high sequence homology between Mfn1 and Mfn2 and our pharmacophore model hypothesis for Mfn2 activation based on conserved residues that regulate HR1-HR2 interactions (Figure S1). Furthermore, it is consistent with the work by Nunnari and colleagues that suggests heterotypic complexes possess greater efficiency in fusion compared to homotypic Mfn1 or Mfn2 complexes (Hoppins *et al.*, 2011) Strikingly, MASM7 emerged as a potent inducer of fusion, achieving an EC_50_ approximately four times lower than 367-384 Gly (MASM7 EC_50_ = 75 nM; 367-384 Gly EC_50_ = 340 nM) (Figure 3D). Furthermore, while MASM7 markedly increased mitochondrial aspect ratio in MFN2 KO MEFs transduced with WT Mfn2, MASM7 did not increase mitochondrial aspect ratio in MFN2 KO MEFs transduced with D725A/L727A Mfn2, underscoring the specificity of MASM7 (Figures 3E and 3F). Taken together, the specificity of the MASM7-mitofusin interaction is evidenced by mutagenesis using NMR, FRET and mitochondrial fusion studies (Figures 2E, 2F, 2H and 3F).

**Figure 3.**
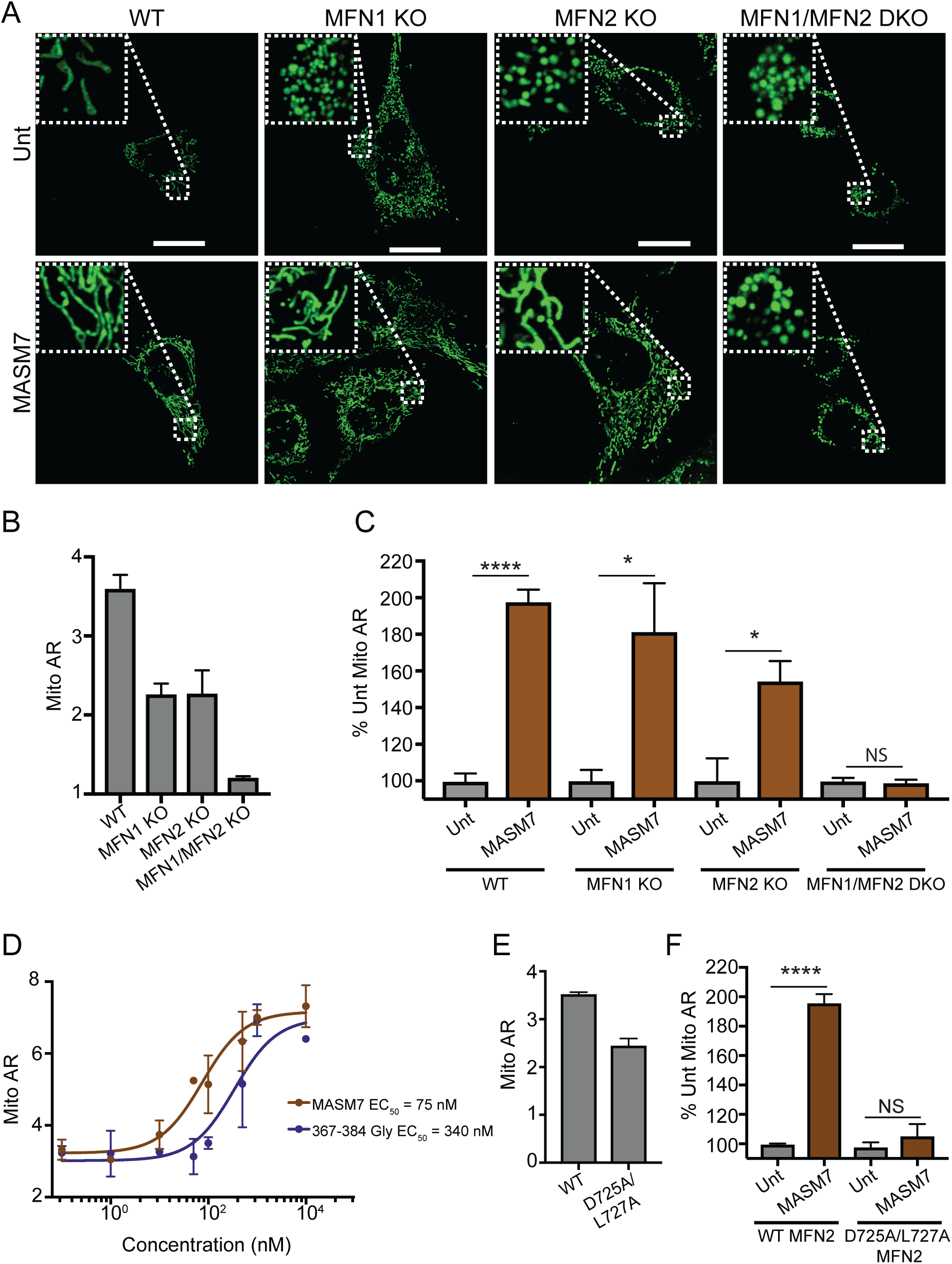
MASM7 treatment induces mitochondrial fusion in WT and mutant MEFs. (A) MASM7 induced mitochondrial fusion in WT, MFN1 KO and MFN2 KO MEFs, but not in MFN1/MFN2 DKO MEFs. Cells were treated with MASM7 (1 μM, 2 h). Mitotracker green stains mitochondria. Scale bar 20 μm. Zoomed images are 5x. (B) Quantification of mitochondrial aspect ratio (Mito AR) in untreated WT, MFN1 KO, MFN2 KO MEFs and MFN1/MFN2 DKO MEFs. Error bars represent mean ± SD, *n* = 3. >200 mitochondria were scored per condition. (C) Relative increase of Mito AR in the indicated cells upon MASM7 treatment. Error bars represent mean ± SD, *n* = 3. >200 mitochondria were scored per condition. (D) MASM7 and 367-384 Gly concentration-responsively increased Mito AR in WT MEFs. Cells were treated with MASM7 or 367-384 Gly at the indicated concentrations for 2 h. Error bars represent mean ± SEM, *n* = 3. (E) Quantification of mitochondrial aspect ratio (Mito AR) in untreated MFN2 KO MEFs reconstituted with WT MFN2 or D725A/L727A MFN2. Error bars represent mean ± SD, n = 2. >200 mitochondria were scored per condition. (F) Relative increase of Mito AR in MFN2 KO MEFs reconstituted with WT MFN2 or D725A/L727A MFN2 upon MASM7 treatment. Error bars represent mean ± SD, n = 2. >200 mitochondria were scored per condition.

## Conclusions

In conclusion, we have identified the first direct small molecule activator of mitofusins using a pharmacophore model small-molecule screen based on the interaction of HR1-HR2 domains of Mfn2. Our studies support the concept that disruption of HR1-HR2 interactions is critical for mitochondrial tethering and fusion to take place. MASM7 promptly and robustly promotes mitochondrial fusion in a mitofusin-dependent manner. The ability to directly activate mitofusins with small molecules provides new opportunities to elucidate the precise roles of these proteins in mitochondrial dynamics as well as the role of mitochondrial fusion in fundamental physiological processes. As importantly, the discovery of MASM7 raises the possibility of novel therapeutics. These may be applied to orphan syndromes such as Charcot-Marie-Tooth disease type 2a resulting from mutation in one Mfn2 allele, as we have recently shown (Rocha *et al.,* 2018). Moreover, direct mitofusin activation may also present therapeutic opportunities for common acquired syndromes such as heart failure, neurodegeneration, cancer, and diabetes in which functional defects in mitochondrial fusion contribute to pathogenesis (Shirihai *et al.*, 2015; Schrepfer *et al.*, 2016).

## Acknowledgements

We thank Louis Hodgson for his help with FRET studies and David Chan for providing MEF cell lines. Studies were supported by NIH grants 1R01CA178394 (E.G.) and 5R01HL128071 (R.N.K.). Support was also provided by NIH awards 1S10OD016305, P30CA013330 and American Heart Association Collaborative Science Award (E.G. and R.N.K).

## Methods

### Structural Model of Mfn2

The structural model of Mfn2 was calculated based on the I-TASSER (Iterative Threading ASSEmbly Refinement) hierarchical approach to protein structure as we previously described (Yang *et al.*, 2015; Franco *et al.*, 2016). Top solutions derived from I-TASSER approach were based on the bacterial dynamin-like protein structure in the PDB (ID: 2J69). Energy minimization and analysis of the top-ranked structure was performed with MAESTRO tools (Maestro, v10.5, Schrödinger, LLC). PyMOL (The PyMOL Molecular Graphics System. Version 1.8; Schrödinger, LLC) was used for preparing the figures.

### Ligand preparation library

eMolecules (www.emolecules.com) library of purchasable compounds was converted to 3D structures using LIGPREP (LigPrep, version 3.8, Schrödinger, LLC) and EPIK (Epik, version 3.6, Schrödinger, LLC) generating an *in silico* library of approximately 13.8 million compounds containing compounds with different ionization state at pH 7.0 ± 2.0, stereochemistry and tautomeric form, excluding potential Pan Assay Interference Compounds (PAINS) using PAINS definitions included in Canvas (Baell *et al.*, 2014). Conformation analysis of ligands was calculated using the OPLS3 force field.

### 3D Pharmacophore model generation and screen

Phase (Phase, version 4.6, Schrödinger, LLC) module was used to generate a pharmacophore hypothesis and a 3D pharmacophore screen (Dixon *et al.*, 2006). The coordinates of the HR1 helix residues Val372, Met376 and His380 from the structural model of Mfn2 were used to assign pharmacophore points in 3D coordinates. Pharmacophore hypothesis included 5 features as defined in Phase for 3 hydrophobic groups to mimic the sidechain residues of Val273 and Met376 and an aromatic ring with a hydrogen-bond donor to mimic the sidechain of His380. The pharmacophore screen used the *in silico* library of compounds prepared from the eMolecules library in pre-existing conformations with the requirement to satisfy at least 4 out of the 5 pharmacophore features of the hypothesis. The top 1000 compounds ranked based on the Phase Score were selected for further visual analysis and clustered for diversity using dendritic fingerprints in Canvas (Sastry *et al.*, 2010). Physicochemical and AMDET properties including Lipinski rules, permeability, logP, metabolic liabilities and hERG inhibition were evaluated using QikProp (QikProp, version 3.4, Schrödinger, LLC, New York, NY, 2011). The highest 8 ranked compounds and the 10 most diverse compounds yielded selected molecules for experimental validation. MASM7 was checked for potential Pan Assay Interference Compounds (PAINS) (Baell *et al.*, 2014) and has not been reported as a hit in previous screens in Pubchem database.

### Compounds

Pro-fusion 367-384 Gly peptide, QIAEVRGIMDSLHMAARGGYGRKKRRQRRR was synthesized, purified at >95% purity by Life Technologies. MASM7 compound was obtained from Enamine (cat. # EN300-396282). Screened MASMs were purchased from Enamine, ChemBridge and ChemDiv. All compounds were >95% pure, dissolved in DMSO to prepare a 10 mM stock solution.

### Cell lines

Cell lines were purchased from ATCC. MEFs (WT, Mfn1 KO, Mfn2 KO and Mfn1/Mfn2 DKO) were also provided by David Chan’s laboratory. HEK 293T cells were provided by Louis Hodgson’s laboratory. All cells maintained in DMEM (Life Technologies) supplemented with 10% FBS, 100 U ml^−1^ penicillin/streptomycin and 2 mM L-glutamine.

### Molecular cloning

To generate FRET-based Mfn2 biosensor construct, we utilized pcDNA3.1-Flag-Mfn2 (5’Flag epitope-tagged human Mfn2) as backbone, and fused mCerulean (cyan fluorescent protein; CFP) and mVenus (yellow fluorescent protein; YFP) onto the 5′ and 3’ ends respectively of the Mfn2 open reading frame (Nagai *et al*., 2002; Rizzo *et al*., 2004; Yun *et al*., 2013). The construct was confirmed by sequencing. FRET-based Mfn2 biosensor mutant D725A/L727A was generated by site-directed mutagenesis of the wild type probe.

### FRET assay

HEK 293T cells (5 × 10^5^ cells/well) were seeded in a pre-coated with poly L-lysine 6-well clear bottom plate. Cells were transfected with the total amount of 2.5 μg of DNA per well for the indicated Mfn2 FRET constructs using Lipofectamine® 3000 Reagent (Invitrogen) according to the manufacturer’s protocol. The following day, transfected cells were transferred to a pre-coated with poly L-lysine 96-well black plate (2 × 10^4^ cells/well). The following day, cell media was replaced with FluoroBrite™ DMEM (Invitrogen) supplememented with 10% FBS, 100 U ml^−1^ penicillin/streptomycin, 2 mM L-glutamine and 1 mM GTPγS 15 min prior treatments with MASMs. GTPγS (Sigma Aldrich) was used to inhibit GTPase activity of Mfn2. Subsequently, cells were treated for 2 h with MASMs or vehicle (0.1% DMSO). Dilutions of MASMs were performed using a TECAN D300e Digital Dispenser from 10 mM stock solutions. Fluorescence intensity was detected by a M100 microplate reader (TECAN), filters that were used for the FRET studies: donor (mCerulean) Ex: 436 nm/Em: 480 nm, acceptor (mVenus) Ex: 436 nm/Em: 535 nm. Acceptor/Donor ratio referred to the text and figures correspond to the ratio of fluorescence intensities (relative fluorescence units: RFU) detected by the acceptor filter/donor filter. Non-transfected cells were used to subtract background fluorescence from each of the aforementioned filters that were used in the study.

### NMR samples and experiments

Mfn2 residues 678-757 corresponding to the HR2 domain were cloned into a pET-28 vector and transformed into BL21(DE3) CodonPlus (DE3)-RIPL *E. coli* cells. The double mutant D725A/L727A construct was also prepared by site-directed-mutagenesis. Cells were grown at 37°C in 1 L of LB media to an OD_600_ of 0.8, cells were then harvested and resuspended in 1 L of M9 media supplemented with 1.5 gr/L of ^15^N ammonium chloride grown for 45 min at 37°C and induced at 18°C for 16 hours with 1 mM IPTG. MFN2 HR2 domain was purified from bacterial pellets by high-pressure homogenization in lysis buffer (20 mM Tris.HCl pH 7, 250 mM KCl, 25 mM imidazole, and Roche complete EDTA free protease inhibitor cocktail) and ultracentrifuged at 45,000 g for 45 min. The supernatant was applied to pre-equilibrated 1 mL HisPur Ni-NTA Resin washed in lysis buffer and eluted using elution buffer (20 mM Tris.HCl pH 6, 250 mM KCl, 400 mM imidazole). ^15^N-MFN2-HR2 was further purified by size exclusion chromatography (Superdex 75 Increase 10/300 GL column) in gel filtration buffer (20 mM potassium phosphate pH 6, 150 mM KCl). Fractions containing the ^15^N-MFN2-HR2 were confirmed by SDS-PAGE, pooled and concentrated to 50 μM in NMR buffer (20 mM potassium phosphate pH 6, 150 mM KCl, 10% D_2_O) using a 10 KDa cut-off Centricon spin concentrator for NMR analysis. ^1^H-^15^N-HSQC spectra of 50 μM MFN2-HR2 in the presence and absence of MASM7 were recorded on a BRUKER AVANCE IIIHD 600MHz system equipped with a 5mm H/F-TCI CryoProbe at 25°C. Spectra were processed using qMDD (mddnmr v2.0) and NMRPTPE and analyzed using Analysis (CCPNMR) (Vranken *et al.*, 2005; Delaglio *et al.*, 1995).

### Live cell imaging

MEFs were seeded on chamber slides (MatTek Corporation: 35 mm dishes, No. 1.5, 14 mm glass diameter) at ~70% confluence. Cells were treated with MASM7 or 367-384 Gly (1 μM, 2 h). Cells were stained with MitoTracker Green (200 nM, Invitrogen) for 20 min at 37°C. After treatments, media was replaced with FluoroBrite™ DMEM (Invitrogen) supplemented with 10% FBS, 100 U ml^−1^ penicillin/streptomycin, 2 mM L-glutamine and MASM7 or 367-384 Gly (1 μM) prior image acquisition. Images were taken with Leica SP5 inverted confocal microscope. Data were analyzed with Image J software. >200 mitochondria were measured per condition in mitochondrial aspect ratio.

### Immunofluorescence

Transfected HEK 293T cells were washed with once PBS and fixed with 2% PFA for 13 min. Subsequently, cells were blocked, then incubated for 1 h at room temperature with the primary antibody for ATP5B (Abcam; ab128743). After incubation with the primary antibody, cells were washed with PBS and incubated for 1 h at room temperature with the secondary antibody (ThermoFisher Scientific; A11010). Images were taken with Leica SP5 inverted confocal microscope or Zeiss fluorescent microscope. Data were analyzed with Image J software.

**Supplementary Table 1:**
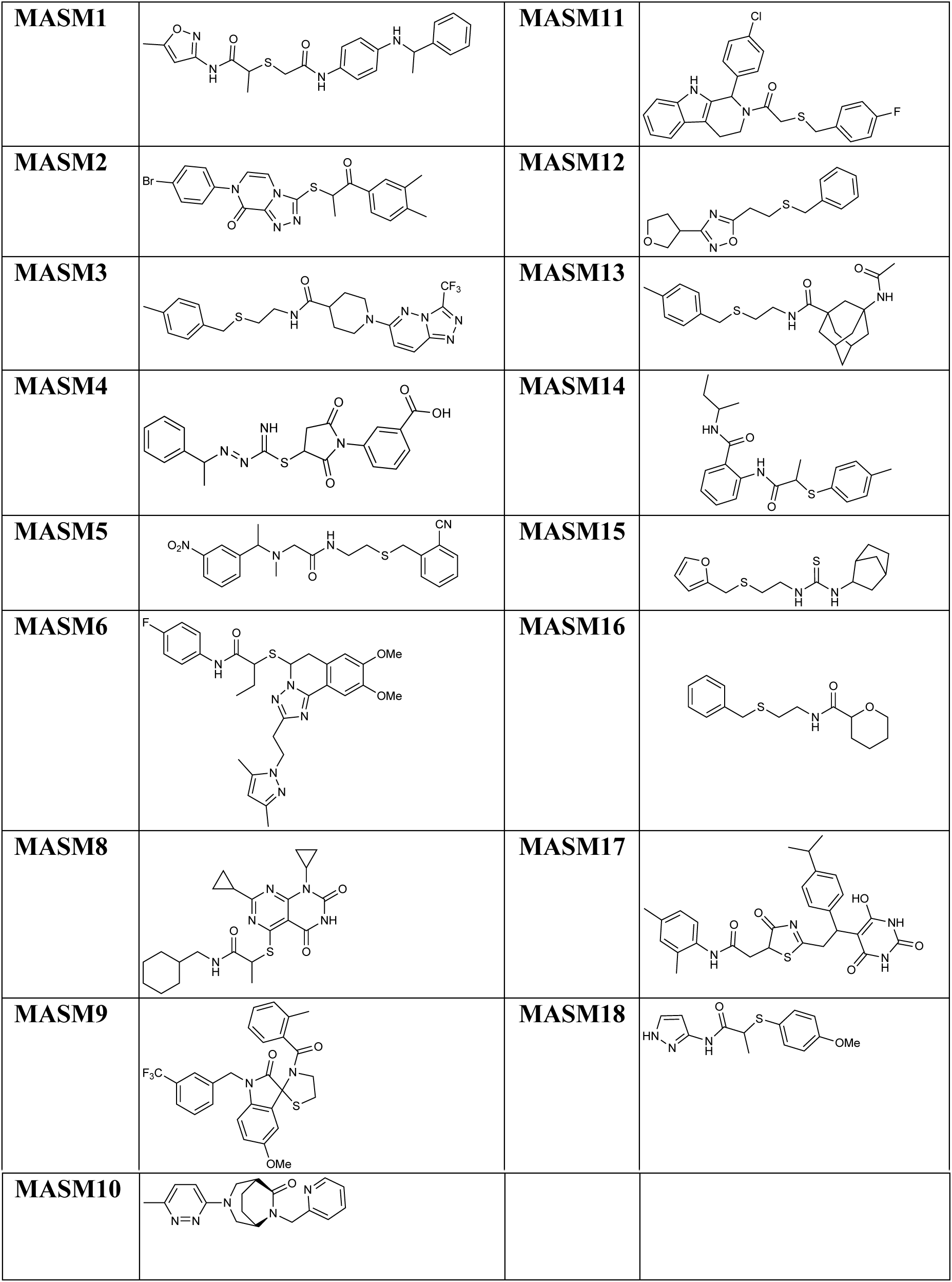
Structures of predicted MASMs.

#### Supplementary Figures

**Figure S1 (related to Figures 1 and 2).**
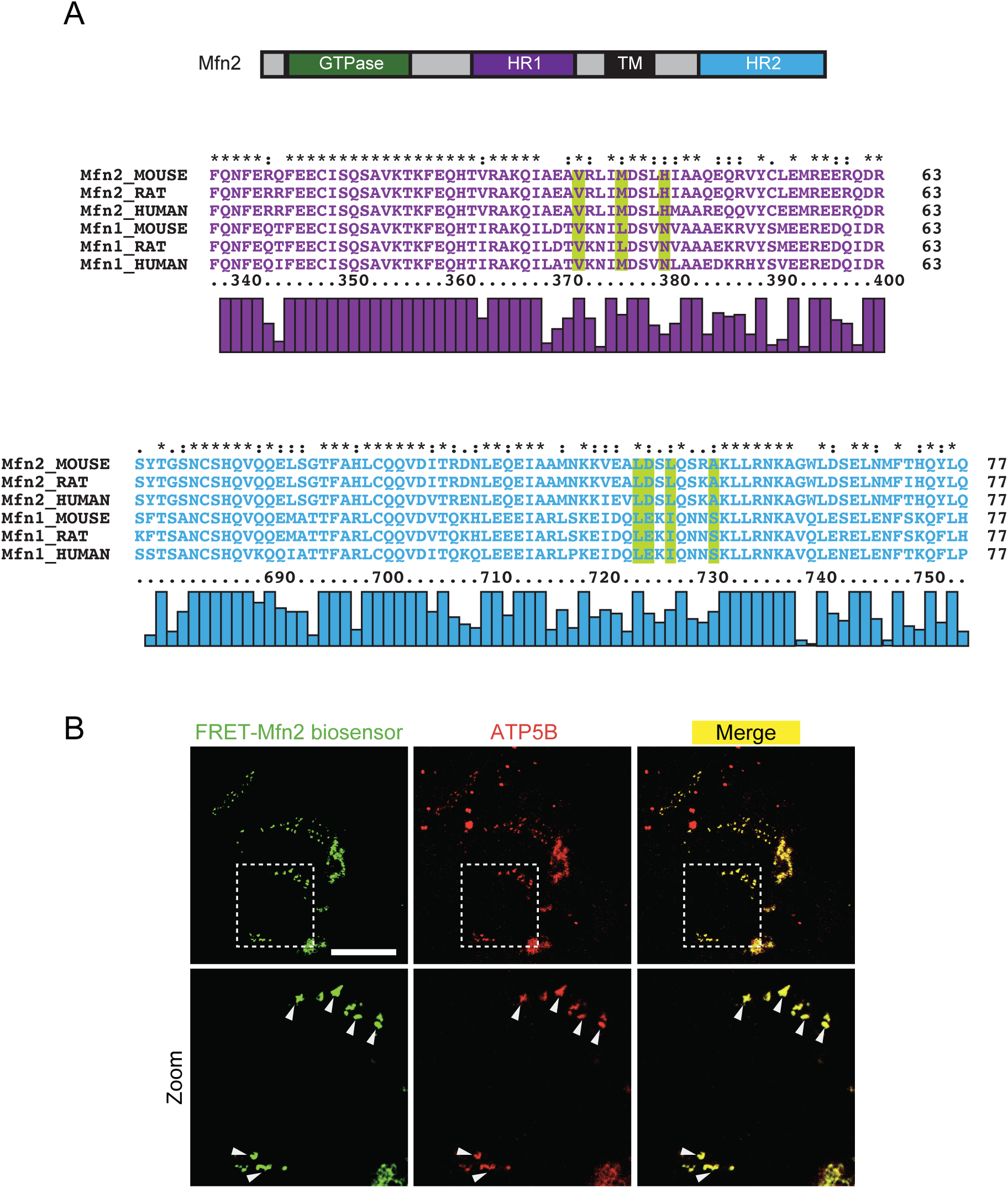
(A) HR1 and HR2 domains in human Mfn1 and Mfn2 shares high sequence homology with mouse and rat Mfn1 and Mfn2. HR1 and HR2 residues used in our pharmacophore hypothesis are highlighted in yellow. (B) FRET-based Mfn2 biosensor co-localizes with ATP5B in mitochondria in HEK 293T cells. Scale bar 20 μm. Zoomed images are 3x.

**Figure S2 (related to Figure 2).**
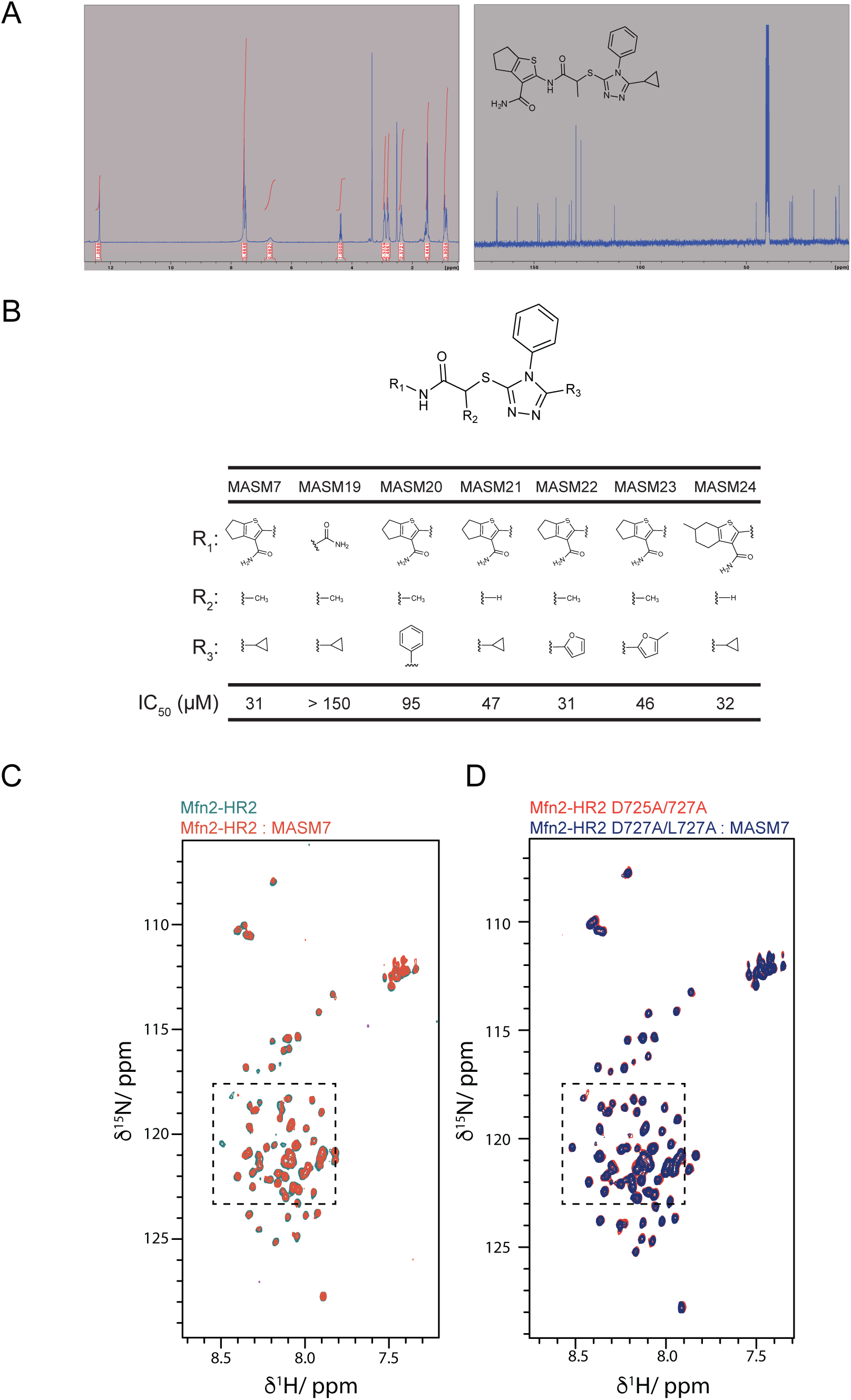
(A) Spectral data of MASM7. ^1^H(left) and ^13^C (right) NMR. (B) Structure-activity relationship of MASM7 scaffold. Potency of MASMs compounds with variable R_1_, R_2_, R_3_ groups were evaluated using WT FRET-based biosensor. HEK 293T cells were treated with MASMs for 2 h. (C) Comparison of the ^1^H-^15^N HSQC spectra of Mfn2-HR2 (50 μM) with (red cross-peaks) and without (green cross-peaks) MASM7 (200 μM). Zoomed region as shown in Figure 2E is highlighted in black box showing overlaid spectra of Mfn2-HR2 with and without MASM7 that demonstrates examples of the observed line broadening and chemical shifts upon addition of MASM7 as highlighted with blue boxes. (D) Comparison of the ^1^H-^15^N HSQC spectra of Mfn2-HR2 D725A/L727A mutant (50 μM) with (blue cross-peaks) and without (red cross-peaks) MASM7 (200 μM). Zoomed region as shown in Figure 2F is highlighted in black box showing overlaid spectra of Mfn2-HR2 D725A/L727A mutant with and without MASM7 that demonstrates loss of effects of MASM7 on broadening and chemical shifts.

